# Dynamic remodeling of hybrid BBB-fenestrated capillaries in the hypothalamic arcuate nucleus

**DOI:** 10.64898/2026.11.15.766735

**Authors:** Aviv Halfon, Yair Oster, Yoel Yeretz-Peretz, Miriam K. Minsk, Doron Kleiman, Batia Bell, Esther Sasson, Shira Anzi, Rachel Ben-Haroush Schyr, Elizabeth E. Crouch, Stefan Liebner, Danny Ben-Zvi, Ayal Ben-Zvi

**Author notes:** Correspondence: Ayal Ben-Zvi -; Danny Ben-Zvi -, Department of Developmental Biology and Cancer Research, The Institute for Medical Research Israel-Canada, Faculty of Medicine, Hebrew University of Jerusalem, 91120 Jerusalem, Israel.

## Abstract

The hypothalamic arcuate nucleus (Arc) is a central regulator of metabolism. We show that capillaries with sparse fenestrae enable direct and rapid hormone transport into the Arc. Arc fenestrae are virtually absent during pregnancy and obesity, but reform postpartum and upon weight-loss. Fenestrated Arc endothelial cells display a unique hybrid BBB-permeable structural and molecular phenotype, which can be regulated by Wnt signaling. The Arc endothelium exhibits functional plasticity that may affect hypothalamic sensing of systemic metabolic signals.

## Main

The Arcuate Nucleus of the hypothalamus (Arc) is a critical metabolic hub, integrating neuronal and systemic hormonal signals such as leptin, GLP-1, and ghrelin to regulate systemic metabolism and feeding behavior^1–3^. The Arc vasculature is considered to possess canonical blood-brain barrier (BBB) features, including restrictive tight junctions and lack of fenestrae: 70-100nm pores spanning the endothelial cells (EC) that enable non-selective rapid transport of molecules. In contrast, highly permeable, fenestrated non-BBB vessels are found at the median eminence (ME), located adjacent and ventral to the Arc at the base of the hypothalamus, facilitating hypothalamic hormone secretion^4–5^.

This vasculature architecture implies that hormones enter the Arc via indirect or slow pathways, such as receptor-mediated BBB transcytosis or transport via the cerebrospinal fluid^6–11^ (Extended Data Fig. 1). However, it was shown that Arc neurons respond to systemic ghrelin administration within less than a minute^7^, implying the potential existence of direct and fast communication avenues between circulating hormones and Arc neurons. Indeed, earlier evidence supported by electron microscopy (EM) and by Plvap staining, identified existence of fenestrated Arc vasculature^6,13,14^.

Using transmission electron microscopy (TEM) and direct super-resolution microscopy (dSTORM) capturing topology of the fenestra structural protein Plvap, we confirmed earlier reports^6,13,14^ and describe sparse fenestration in 30±10% of Arc capillaries in male mice fed *ad-libitum* (Fig. 1a-f). High-resolution imaging approaches are critical for quantifying Arc fenestration since the resolution of confocal light microscopy does not exceeds 200nm, much larger than a single fenestra (Fig. 1d, Extended Data Fig.2a-c). Fenestrated vessels were predominantly located in the ventro-medial Arc neighboring the ME^13^, but were also found in lateral and dorsal Arc regions. In reference, all ME capillaries were fenestrated/Plvap^+^, and all cortical capillaries were non-fenestrated/Plvap^-^ (Fig. 1b,e). Arc vessels displayed intermittent fenestration/Plvap-clusteres along the endothelial circumference in contrast with the dense and continuous ME arrangement (Fig. 1c,f). Both ME and Arc capillaries had indistinguishable fenestrae diameter, but Plvap signal/cluster was higher at the ME (Extended Data Fig.2e,d).

Functional tracer assays confirmed that Arc fenestrae facilitate rapid transport of large molecules from the blood into the Arc parenchyma. One minute after intravenous (IV) injection, 10 kDa dextran and 20 kDa leptin were detected via dSTORM exclusively adjacent to Plvap⁺ clusters (Fig. 1g-k). One-minute long permeability challenges were designed to reduce the contribution of transport via indirect pathways. Extravasation from fenestrated ME capillaries was more extensive than that of fenestrated Arc capillaries, and non-fenestrated Arc capillaries showed nearly no extravasation (Fig. 1h,j, capillaries circumference delineated with either Plvap or Glut1, Extended Data Fig.2f). These findings provide functional evidence that Arc fenestrae act as rapid, non-selective gates for large molecules and peptide hormones.

**Figure 1.**
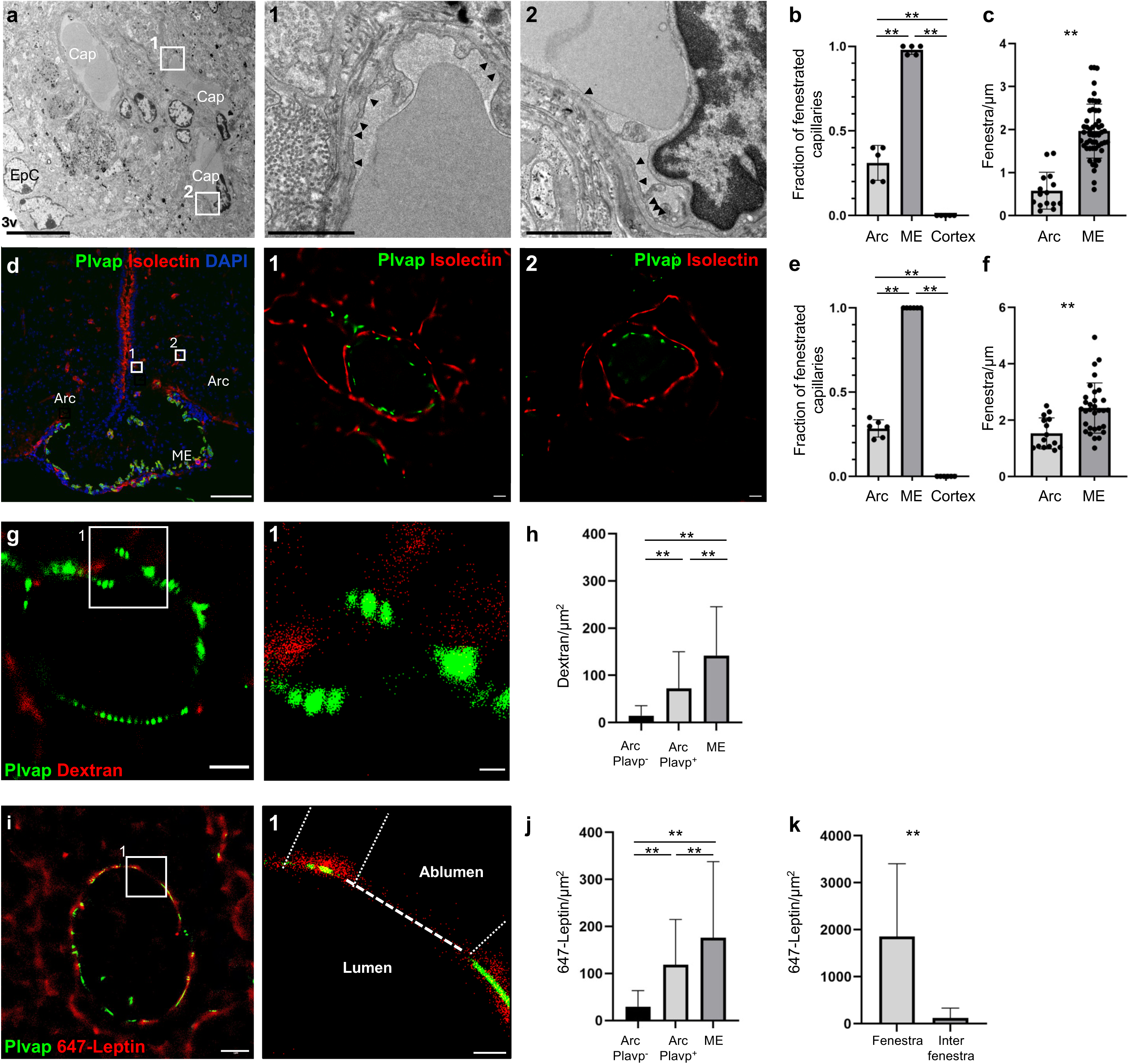
Abundance of Arcuate Nucleus fenestrated vessels and fenestra function facilitating rapid non-selective transport of large molecules. **a**, Transmission electron microscopy (TEM) image of the Arcuate Nucleus (Arc) region adjacent to the third ventricle (3V). Capillaries (Cap) and ependyma cells (EpC) are labeled. Scale bar: 10 μm. Insets (1) and (2) show higher magnification views of the capillary endothelium; black arrow heads indicate endothelial fenestrations. Scale bar: 1 μm. **b,** Fraction of fenestrated vessels within the Arc, ME and Cortex (TEM analysis). **c,** Fenestra density along the circumference of capillaries (TEM analysis). **d,** Confocal imaging of the Arc/median eminence (ME) region. Scale bar: 100 μm. Insets (1) and (2) show higher magnification views of individual vessels using dSTORM imaging of Arc vasculature. Scale bar: 1 μm. **e,** Fraction of fenestrated vessels (dSTORM analysis) within the Arc, ME and Cortex. **f,** Fenestra density along the circumference of capillaries (dSTORM analysis). **g,** dSTORM-based tracer assay demonstrating 10 kDa dextran passage through Plvap^+^ Arc capillaries one-minute after IV injection. **h,** Quantification of dextran signals in the immediate space abluminal to the circumference of capillaries (delineated by Plvap signals, or Glut1 signals, Extended Data Fig.2f), across Plvap^-^ and Plvap^+^ ARC capillaries, and through the ME vesseles (n=39, 52, 14 fenestrae). **i,** dSTORM-based tracer assay demonstrating 647-leptin passage through Plvap^+^ Arc capillaries one-minute after IV injection. Scale bar in g,i=1mm and 0.2mm in magnifications. **j,** Quantification of 647-leptin signals across Plvap^-^ and Plvap^+^ capillaries and through the ME (n=28, 58, 62 fenestrae). **k,** 647-leptin signals abluminal to fenestra domains (Plvap clusters) compared to inter-fenestra regions of Plvap^+^ capillaries (n=100, 100 fenestrae). **p < 0.01, 1-way ANOVA with Tukey post-hoc test (b,e,h,j) and Student’s T-test (c,f,k). Each data point in (b,e) represents a single mouse (20-30 capillary profiles per mouse); Each data point in (c,f) represents a capillary cross-section, Data was collected from 5 mice.

The Arc modulates its responsiveness to hormonal signaling during pregnancy and in obesity^15–17^. We hypothesized that vascular permeability might also change in these conditions, affecting the dynamics of Arc responsiveness to hormonal signals. This corresponds to an earlier finding that peripheral circulating leptin access into the brain is completely suppressed during pregnancy, albeit providing no underlying mechanisms^10^. Indeed, we observed a significant reduction in Arc fenestrations from 30%±10% of vessels in virgin female mice to just 3%±5% (Fig. 2a-c, e-f) in pregnant mice at E16.5, at hyperleptinemia peak^10^. Fenestrae density in remaining fenestrated Arc vessels was also sharply reduced (Fig. 2d, Extended Data Fig. 3a), as was Plvap density per cluster (Extended Data Fig. 3b). These structural changes translated into a significant diminished leptin influx (Fig. 2e,g, Extended Data Fig. 3c). Fenestrae abundance and leptin influx remained significantly lower than in virgin mice three weeks post-partum (P21), and normalized only at P56 (Fig. 2c-d, f-g, Extended Data Fig. 3c).

Diet-induced obesity similarly remodeled the Arc vasculature (Fig. 2h-k). Male mice were fed a high-fat high-sucrose (HFHS) diet for 12 weeks to induce obesity, followed by a normal chow diet for 4 weeks to induce weight loss, or 4 more weeks of HFHS to sustain obesity. Age-matched male mice maintained on a normal chow served as lean controls (see experimental design, Extended Data Fig. 4a). Body weight increased for 12 weeks and kept increasing for 4 weeks in the obese group, while normalizing to levels of lean mice in the weight-loss group (Fig. 2h). Non-fasting glucose measurements demonstrated hyperglycemia in the obese mice, which was normalized in the weight-loss group (Fig. 2i). Plasma levels of lipids and liver enzymes were mostly reduced in the weight-loss group compared to obese mice (Extended Data Fig. 4b-c).

The fraction of fenestrated vessels in the Arc fell to 10%±3% in mice fed a HFHS diet for 16 weeks, which was associated with reduced influx of leptin (Fig. 2j-k, Extended Data Fig. 4d). 22%±7% of the Arc vessels were fenestrated in mice that had a 4-week weight loss period. Obesity is thus associated with a reduction in structural and functional Arc permeability, which are partially reversible upon weight loss.

A similar trend was observed in the ME in pregnancy and obesity, yet the magnitute of fenestrae reduction was much smaller, retaining about 90% of the ME vessels fenestrated (Extended Data Fig. 5a-d). The structural changes in the ME did not translate into significantly changes in leptin influx, tested in pregnancy (Extended Data Fig. 5e). Altogether, dynamic modulation of Arc vasculature during obesity and pregnancy may change hormonal signaling dynamics in the Arc by limiting the rapid access of systemic signals to Arc neurons^7^.

**Figure 2.**
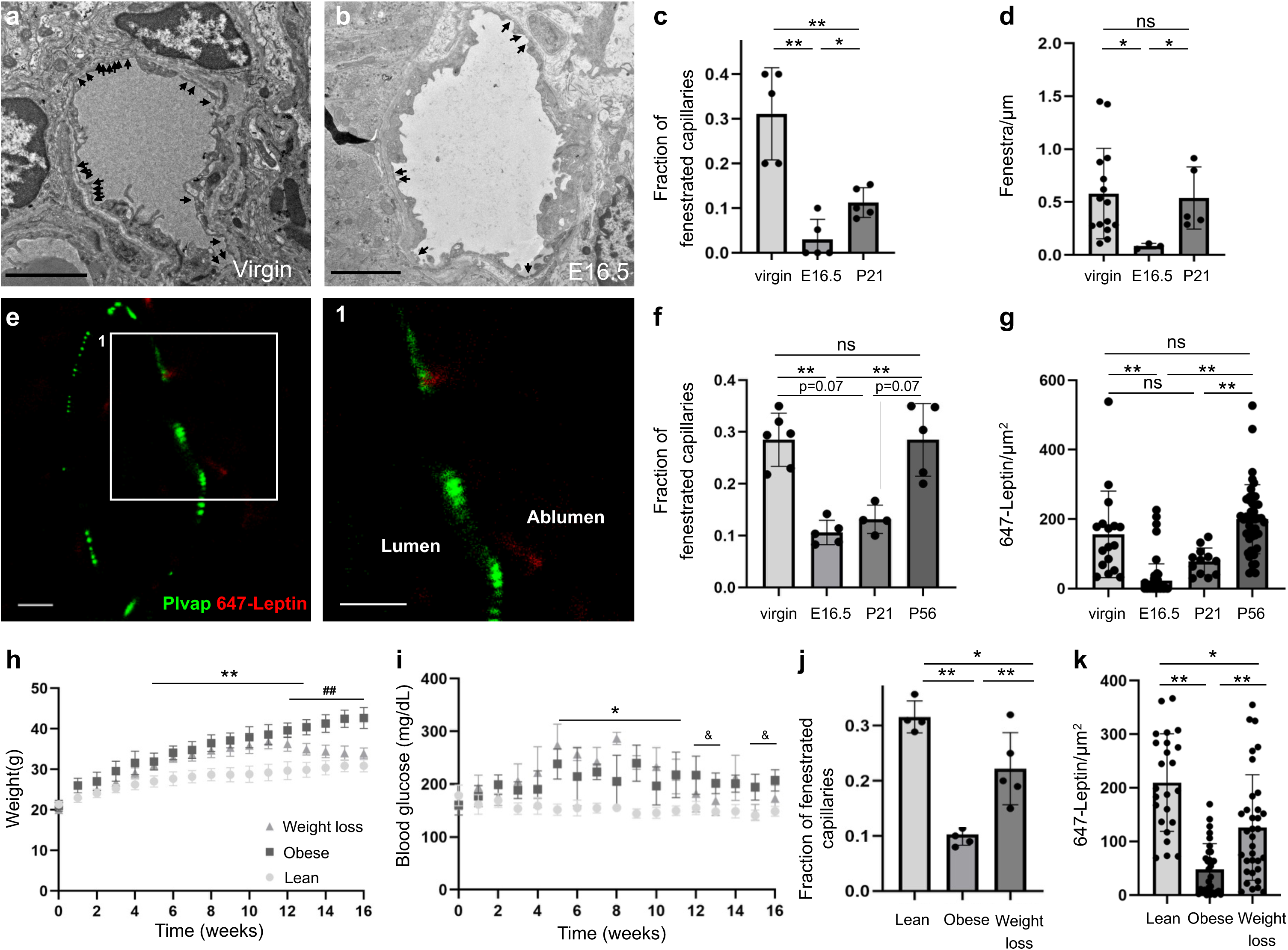
Remodeling of Arc fenestrations during pregnancy and diet-induced obesity. a-b, Representative TEM images of fenestrated capillaries in the Arc of a virgin female mouse (a), and pregnant mouse at E16.5 (b) demonstrating reduced fenestration (arrows) in pregnancy. Scale bar 1μm. **c,** Fraction of fenestrated Arc capillaries in virgin mice, pregnant mice at E16.5 and at P21 (TEM analysis, n=5 mice). **d,** Fenestra density along the circumference of capillaries (TEM analysis) in virgin mice, pregnant mice at E16.5 and at P21. Sample size differed between groups due to the reduced number of fenestrated capillaries in pregnant and postpartum mice (Virgin: n=15 capillaries collected from 5 mice; E16: n=4 capillaries collected from 5 mice; Postpartum: n=6 capillaries collected from 5 mice). **e,** Representative dSTORM image of Arc vessel during pregnancy (E16.5) one minute after IV injection of labeled 647-leptin. scale bar 1μm. Inset: 647-leptin in the abluminal space of a Plvap^+^ capillary. Scale bars: 1mm. **f,** Fraction of fenestrated Arc capillaries in virgin mice, pregnant mice at E16.5, P21 and P56 (dSORTM, n=5 mice). **g,** Quantification of labeled 647-leptin signal in the immediate space abluminal to the circumference of capillaries (delineated by Plvap signals) in the Arc (n≈20 capillaries collected from 5 mice per group). **h-i,** Body weight (h) and blood glucose (i) in obese, lean and weight-loss male mice for 16 weeks (n=5 mice per group). **j,** Fraction of Plvap^+^ (fenestrated) capillaries in the Arc of lean, obese and weight-loss mice (n=5 mice per group). **k,** Density of 647-leptin in the immediate space abluminal to the circumference of capillaries (delineated by Plvap signals) of Plvap^+^ Arc capillaries one minute after IV injection (n≈20 capillaries collected from 5 mice). *p<0.05, **p<0.01 in a Tukey post-hoc test following ANOVA (c,d,f,g,j,k). In h,i:, *p<0.05, **p<0.01 obese, weight loss vs. lean ## p<0.01 obese vs. lean, weight loss. & p<0.05 obese vs. lean; lean vs. weight loss not significant in a Tukey post-hoc test following ANOVA.

Molecular analyses revealed that fenestrated Arc endothelial cells (ECs) display a unique BBB-permeable hybrid phenotype. dSTORM imaging demonstrated that Plvap⁺ Arc capillaries atypically co-express the canonical BBB marker Slc2a1 (Glut1) (Fig. 3a). Analysis of single-cell RNA sequencing (scRNAseq) published datasets^18,19^ from human and mouse hypothalami confirmed existence of a hybrid plvap^+^slc2a1^+^ population of ECs (Fig. 3b,c). Whole transcriptome analysis of BBB-associated gene signatures^20,21^ showed an intermediate BBB score^22^ of these cells in both humans and mice (Fig. 3d,e, Extended Data Fig. 6a,b), and co-expression of *PLVAP* with central BBB-associated genes-such as *MFSD2A* and *CLDN* in both species (Extended Data Fig. 6c,d). Our structural, functional, and histological analyses, together with the unique transcriptomic signature support the existence of a partially-fenestrated BBB EC type in the Arc.

The VEGF pathway positively regulates capillary fenestrations inside and outside the CNS^14,23–28^, and in particular in the Arc-ME border^6,14^. The Wnt signaling pathway is a master regulator of BBB formation and maintenance, and negatively regulates capillary fenestrations in circumventricular organs (CVOs)^28–34^. scRNAseq analysis from human and mouse hypothalami showed no significant difference in gene expression associated with VEGF signaling (not shown). However, the expression of the Wnt target gene and transcriptional factor *Lef1* was high in canonical BBB cells (slc2a1^+^plvap^-^ ) and in the hybrid population (slc2a1^+^plvap^+^, potentially Arc), and lowest in the non-BBB cells (slc2a1^-^plvap^+^, potentially ME, Fig. 3f,g). Nuclear localization of Lef1 provides a functional proxy for the Wnt pathway output. We used single molecule localization microscopy (SMLM-dSTORM) to image immunostained Lef1 in nuclei of cortical, ME and Arc capillaries in *ad-libitum* fed mice. Lef1 was localized to the nucleus of cortical BBB Slc2a1^+^ ECs and was negligible in Plvap^+^ ME ECs. In the Arc, Lef1 levels was variable, and lower compared to the cortex but higher compared to the ME (Fig. 3h, Extended Data Fig. 7). Thus we hypothesized that lower Wnt activation might be permissive for fenestrea formation. We utilized transgenic mice with an inducible constitutively active β-catenin in ECs to test if high Wnt signaling affects Arc fenestration. Three weeks after induction, Arc fenestrations were completely abolished (Fig. 3i-k), mirroring the effect seen in subfornical CVO^34^ and the ME (Extended Data Fig. 8). These results indicate that Wnt signaling can act as a potent negative regulator of Arc fenestrae and has the potential to drive vascular remodeling observed during metabolic challenges.

**Figure 3.**
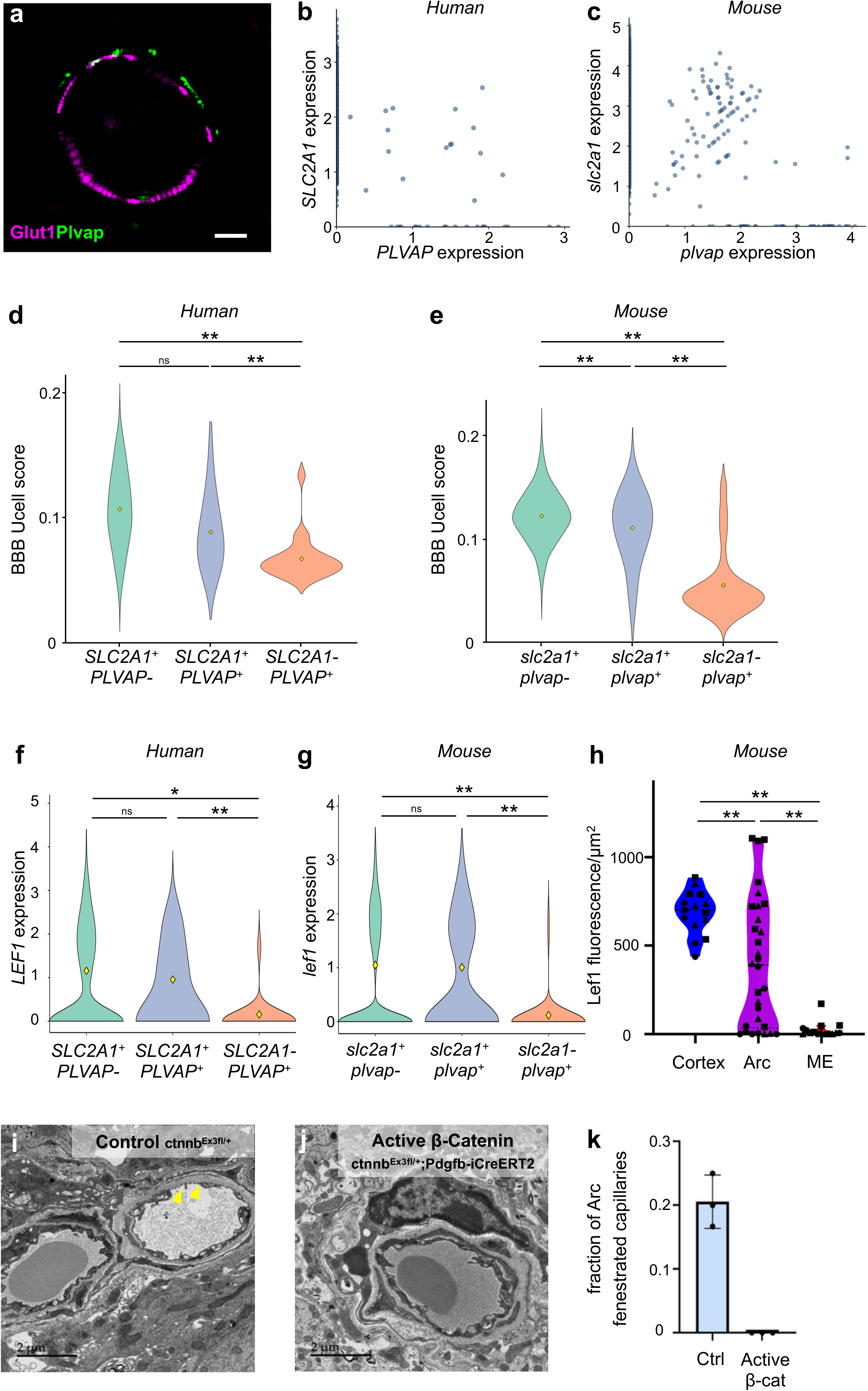
Molecular characterization of fenestrated endothelial cells of the Arc and their Wnt responsivness. a, dSTORM image showing co-localization of Plvap (green) and Slc2a1 (magenta) in an Arc vessel. Scale bar: 1 μm. **b-c,** Single-cell RNA-seq analysis of human (b) and mouse (c) hypothalamic endothelial cells identify a canonical BBB slc2a1⁺/plvap^-^ population, a smaller population of canonical fenestrated slc2a1^-^/plvap⁺ cells, and a third hybrid population of slc2a1^+^/plvap⁺ cells. **d-e,** ‘BBB score’ analyses of the three EC populations of human (d) and mouse (e) hypothalamic scRNAseq data sets. **f-g,** Expression levels of the Wnt downstream transcription factor Lef1 across EC subpopulations identified in human (f) and mouse (g) hypothalamic scRNAseq datasets. **h,** dSTORM imaging analysis of Lef1 nuclear levels in cortex, Arc, and ME capillaries (n=5-10 nuclei of 3 mice per group). **i-j,** Representative TEM images of Arc capillaries in Ctnnb1^Ex3fl/+^ mice^35^ (Control right vessel, i. Left vessel is an arteriole) and Ctnnb1^Ex3fl/+^;Pdgfb-iCre^ERT2^ mice (Active EC b-Catenin, j), 3 weeks after treatment with tamoxifen. Scale bar: 2 μm **k,** Quantification of the fraction of fenestrated vessels in the Arc in control and active β-catenin mice. Each data point represents one mouse (n=20 capillaries for each mouse, 3 mice per group). *p<0.05, **P < 0.01. Statistical comparisons were performed using a linear mixed effects model followed by Tukey’s post-hoc test (d-g) and Kruskal-Wallis test followed by Dunn’s post-hoc test (h)^36,37^.

Our study demonstrates that the Arc vasculature functions as a dynamic gate that includes sparse but functional fenestrae. These fenestrae enable rapid influx of hormones, a structure that is down-regulated during pregnancy and obesity, and re-forms after birth and upon weight loss. Arc fenestraiton might modulate the dynamics of hypothalamic response to hormones during metabolic challenges (illustrated in Extended Data Fig. 9). In these conditions multiple patways mediating hormone expusre might integrate the overall response (see disscusion in Extended text related to main Fig. 2).

Molecularly, these fenestrated blood vessels reside in a hybrid BBB-permissive state, that has lower expression of barrier signature genes, and lower Wnt activity. We provided transcriptional indications for the existence of hybrid endothelial cells also in the human hypothalamus, but further studies are required to verify and characterize the ECs of the Arc in humans at the ultrastructural and protein levels.

CNS vasculature is classically divided into fenestrated at CVOs, and BBB vasculature elsewhere. The discovery of a hybrid endothelial phenotype suggests that CNS endothelium is not adequately represented by a binary classification. Intermediate endothelial phenotypes may be particularly relevant at anatomical interfaces between restrictive and permissive vascular territories, such as the Arc-ME boundary.

In conclusion, Wnt-responsive vascular plasticity of the Arc represents a previously underappreciated layer of metabolic control and a potential mechanism for the modulation of hormonal signaling in physiological and pathophysiological states.

## Methods

### Animals

All mice were maintained in the animal facility of the Hebrew University under specific pathogen-free (SPF) conditions. All experimental procedures were performed in accordance with institutional guidelines approved by the Institutional Animal Care and Use Committee (IACUC) at the Hebrew University (protocol MD-23-17309-3). 8-week-old C57BL/6JOlaHsd wild type male and female mice were purchased from Envigo (Rehovot, Israel) and housed under standard conditions with ad libitum access to food and water unless otherwise specified. Pregnant mice were obtained following overnight mating (day of vaginal plug was defined as embryonic day 0.5). Ctnnb1Ex3^fl/ fl^ mice were a kind gift from Prof. Makoto Mark Taketo, (Kyoto University, Japan)^35^, Pdgfb-iCreERT2 mice were a kind gift from Prof Marcus Fruttiger (University College London, London, UK)^38^.

### Diet-induced obesity model

To induce metabolic obesity, male mice were placed on a high-fat high-sucrose diet (HFD) (Envigo TD08811) beginning at 6 weeks of age and maintained on this diet for 12 weeks. Control mice were maintained on normal chow (NC) throughout the experiment. Following the initial 12-week HFD period, a subset of HFD-fed mice was returned to normal chow for an additional 4 weeks to assess the reversibility of diet-induced metabolic alterations. Body weight and non-fasting blood glucose levels were monitored throughout the experimental period. Liver enzymes (ALT, AST, and ALP) and lipids (total cholesterol, LDL, HDL, and triglycerides) were measured with Cobas c111 analyzer (Roche Diagnostics), according to manufacturer’s instructions.

### Tissue preparation

After dissection, brains were fresh frozen in TissueTek OCT (Sakura). Frozen brains were cut at 6-8 µm sections for dSTORM (CM1950, Leica) and for confocal microscopy to produce coronal brain sections.

### Immunofluorescence

Cryosections were post-fixed in 4% paraformaldehyde (PFA, Sigma Aldrich) for 10 min at room temperature (RT), followed by three washes in phosphate-buffered saline (PBS) for 5 min, incubated for 3 h at RT in blocking solution (10% goat serum (GS), 10% horse serum (NHS), and 0.05% Triton X-100 in PBS). Slides were subsequently incubated with primary antibodies diluted in antibody dilution buffer (2.5% GS, 2.5% NHS, 0.05% Triton X-100 in PBS) overnight at 4°C (see antibodies table for details; Table 1). Slides were then washed three times with PBS for 5 min, incubated with secondary antibodies for 1 h at RT and washed again three times with PBS for 5 min. Samples were mounted with freshly made imaging buffer for dSTORM (describe in the dSTORM imaging section), or mounted in Fluoromount G (EMS) for confocal and epifluorescence microscopy.

**Table 1:** Antibodies details.

| Epitope | Host | Catalogue no. | Company | Dilution |
|---|---|---|---|---|
| Meca-32 (Plvap) | Rat | 553849 | BD | 1:200 |
| SLC2A1(GLUT1) | Rabbit | 071401 | Merck | 1:500 |
| Lef1 | Rab | 2286 | Cell Signaling | 1:400 |

**Table 2:** Fluorophores details.

| Fluorophore | Isotype | Catalouge no. | Company | Dilution |
|---|---|---|---|---|
| Alexa Fluor488 | Anti-rat IgG | 712-545-153 | Jackson | 1:500 |
| Alexa Fluor488 | Anti-rabbit IgG | 711-545-152 | Jackson | 1:500 |
| Isolectin 568 | ------------------- | I21412 | ThermoFisher | 1:500 |
| CF568 | Anti-rabbit IgG | 20098 | Biotum | 1:500 |
| Alexa Fluor568 | Anti-rat IgG | A11011 | ThermoFisher | 1:500 |

### Leptin labeling with Alexa Fluor 647

Recombinant mouse leptin (R&D Systems, R98-OB) was fluorescently conjugated to Alexa Fluor 647 NHS ester (Thermo Fisher Scientific, A20006) using standard NHS-ester coupling chemistry as previousaly described^39^. Lyophilized leptin (5 mg) was resuspended in phosphate-buffered saline (PBS, pH 7.4, Ca²⁺/Mg²⁺-free). Alexa Fluor 647 NHS ester was dissolved in anhydrous DMSO to generate a 20 mM stock solution. Leptin was incubated with Alexa Fluor 647 NHS ester at an approximately 1:1 molar ratio for 1 h at room temperature under gentle rotation, protected from light. The reaction was quenched by addition of Tris buffer (final concentration 50 mM, pH 8.0).

Unconjugated dye was removed using Zeba spin desalting columns (7K MWCO, Thermo Fisher Scientific) equilibrated with cold PBS. Samples were passed sequentially through two desalting columns according to the manufacturer’s instructions to ensure efficient removal of free dye.

Protein concentration of the labeled leptin preparation was estimated using a NanoDrop spectrophotometer. Labeled leptin was kept on ice and used immediately for in vivo injections at a dose of 7.5 mg/kg^39^.

### Permeability assay

To assess vascular permeability in the arcuate nucleus (Arc), in vivo tracer experiments were performed using two tracers with distinct biological properties. An inert fluorescent tracer, Dextran (10 kDa), was used to evaluate non-specific vascular permeability, while fluorescently labeled leptin was used as a biologically active hormone tracer. For permeability analysis, a leakage event was defined when tracer signal was detected outside the endothelial boundary, as determined by endothelial markers or vascular ultrastructure. Quantification of vascular leakage was performed by calculating the relative leakage index, defined as tracer signal density (signals per area) within a fixed region of interest at a defined distance from the luminal side of the vessel. This approach allowed assessment of both non-specific permeability using dextran and biologically mediated hormone transport using leptin across Arc vasculature.

| Tracer | Size | Company | Catalog No. |
|---|---|---|---|
| Alexa Fluor647 anionic fixable conjugated dextran | 10kDa | ThermoFisher | D22914 |
| Alexa Fluor 647–conjugated leptin | ~16 kDa | ThermoFisher | A20006 |

Mice were anesthetized with isoflurane (N0115B28C) and injected IV with fluorescent tracers. Mice received either Alexa Fluor 647-conjugated dextran or Alexa Fluor 647-conjugated leptin prepared as described above. Tracers were diluted in sterile PBS. To capture rapid transvascular transport events, tracers were allowed to circulate for 1 minute prior to tissue collection. Following circulation, animals were sacrificed and brains were rapidly dissected and processed as described in the tissue preparation section.

### Transmission electron microscopy (TEM)

Freshly dissected tissue samples were fixed with Karnovsky’s fixative (2% PFA,

2.5% glutaraldehyde in 0.1 M cacodylate buffer, pH = 7.4) for 4 h at RT, followed by a 1:2 dilution of Karnovsky’s fixative in 0.1 M cacodylate buffer overnight at 4 °C. 80 μm thick sections were prepared with a vibratome (VT1000 S, Leica), and ME-Arc regions were manually dissected, these samples were post-fixed in 1% Os04 in 0.1 M cacodylate buffer for 2 h, dehydrated in a graded series of alcohols, and embedded in epoxy resin. Sections of 60–80 nm were cut on an ultramicrotome (Ultracut, Reichert-Jung), contrasted with uranyl acetate and lead citrate, and examined with a Jeol (JEM-1400 PLUS, Japan) electron microscope.

### Fluorescent microscopy

Immunofluorescence images were captured using the following confocal microscopes: Nikon Eclipse Ni, objective ×20 and ×40 with a Nikon C2 camera, and Nis-Elements software, or with Nikon TE-2000, objective ×20, ×40 and ×60 with EZ-C1 software.

### dSTORM imaging

Direct stochastic optical reconstruction microscopy (dSTORM) was performed to achieve nanoscale visualization of vascular structures. Imaging was carried out using a Nikon N-STORM system based on TIRF illumination, enabling a spatial resolution of approximately ∼20 nm through the use of photoswitchable fluorophores. Brain sections (6-8 μm thickness) were mounted on poly-D-lysine coated coverslips (No. 1.5H, Marienfeld-superior, Lauda-Königshofen, Germany). Imaging was performed in freshly prepared dSTORM imaging buffer containing 0 mM Tris (pH 8.0), 10 mM NaCl and 10% (w/v) glucose with an oxygen-scavenging GLOX solution (0.5 mg/ml glucose oxidase (Sigma-Aldrich)), 40 μg/ml catalase (Sigma-Aldrich), 10 mM cysteamine MEA (Sigma-Aldrich), and 1% β mercaptoethanol 61–63. A Nikon Ti-E inverted microscope was used. The N-STORM (Nikon STORM system) was built on TIRF illumination using a 1.49 NA X100 oil immersion objective and an ANDOR DU-897 camera. 488, 568 and 647 nm laser lines were used for activation with cycle repeat of ∼8000 cycles for each channel. Nikon NIS Element software was used for acquisition and analysis; analysis was also performed by NIH ImageJ^40^.

### Super-resolution cluster analysis

Single-molecule localization microscopy (SMLM) datasets obtained from dSTORM imaging were processed using ThunderSTORM (ImageJ) to generate molecular localization lists containing the spatial coordinates of individual detected molecules. To identify Plvap-positive molecular clusters, localization data were analyzed using custom MATLAB scripts (MathWorks)^41,42^. Pairwise distances between localization points were calculated and molecules were grouped into clusters using hierarchical clustering based on spatial proximity. Two localization points were considered part of the same cluster when their distance was below a predefined threshold. The clustering threshold was determined empirically based on ultrastructural measurements of endothelial fenestrae obtained by TEM analysis revealed fenestra diameters ranging between approximately 40–80 nm, and therefore a 50 nm threshold distance was used to define Plvap clusters in super-resolution datasets. This threshold was selected based on calibration analysis comparing fenestra size measured by TEM with Plvap cluster size measured by dSTORM (Extended Data Fig. 10). Cluster spatial distributions were visualized and quantified to assess the structural organization of Plvap-positive domains across experimental conditions.

### Transcriptomics analyses

Published scRNAseq data from mouse and human hypothalami were analyzed^18,19^ using Seurat v5 (https://pubmed.ncbi.nlm.nih.gov/37231261/). Hybrid ECs were specifically validated not to be doublets.

Blood-brain barrier (BBB) module scores were calculated using previously published BBB-associated gene signatures^20,21^, following an approach adapted from Nano et al.^22^. For each endothelial cell class, gene signature scores were calculated using UCell in Seurat. The BBB gene signatures were originally defined using mouse genes; mouse genes were mapped to their human orthologs prior to scoring the human dataset.

Continuous module scores (BBB signature) and log-normalized transcript counts (LEF1/lef1) across endothelial subpopulations were evaluated using hierarchical linear mixed-effects models via lme4 (v1.1-36) and lmerTest (v3.1-3) with post-hoc Tukey’s adjustment^36,37^.

### Statistical analysis

All statistical analyses were performed using GraphPad Prism 8.0.1 (GraphPad Software, San Diego, CA, USA). Data distribution was assessed prior to statistical testing. Comparisons between two groups were performed using either an unpaired two-tailed Student’s *t*-test for normally distributed data or a two-tailed Mann–Whitney *U* test for non-normally distributed data. Comparisons among three or more groups were performed using one-way ANOVA with appropriate post hoc multiple-comparison tests for normally distributed data, or the Kruskal-Wallis test with Dunn’s multiple-comparison correction for non-parametric data. Data are presented as mean ± s.e.m. A *P* value < 0.05 was considered statistically significant. Unless otherwise indicated, each data point represents an individual biological replicate (mouse, blood vessel, or fenestra, as specified in the corresponding figure legend). Sample sizes (*n*) are indicated in each figure or figure legend.

AI assisted editing and illustration (Extended Data Fig. 7a) was done using Google Gemini.

## Acknowledgments

We would like to thank; Danny and Ayal Ben-Zvi groups for scientific inputs, and the electron microscopy unit, especially Mrs. Hava Glickstein for valuable technical support (the intra-departmental core facility at the Hebrew University Medical School).

---

**Extended Data Fig. 1.**
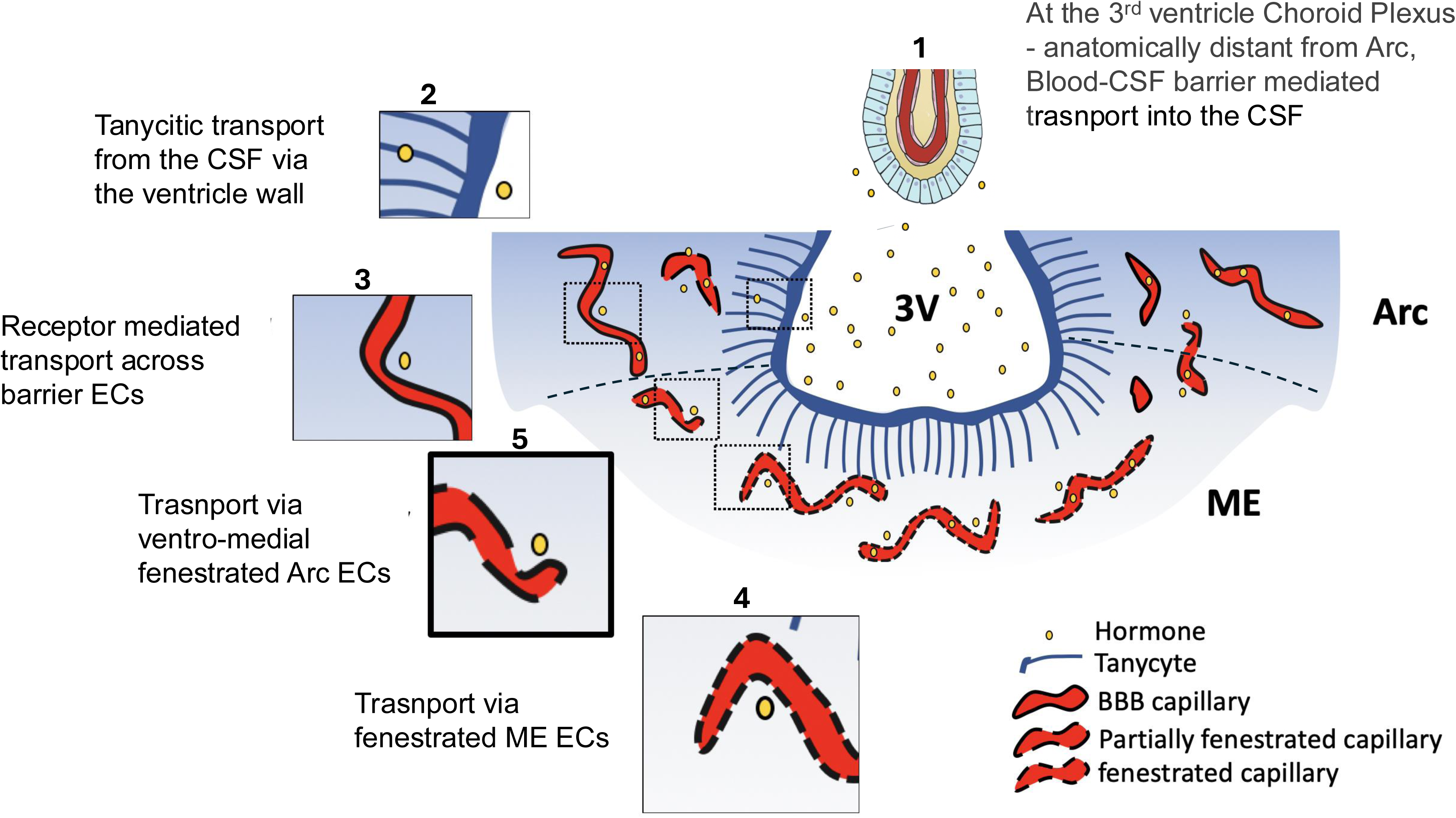
Illustration of indirect pathways for hormones accesses into the Arc. Hormones are suggested to enter the Arc from the blood by several indirect paths: Hormones can enter the cerebrospinal fluid (CSF) through a blood-CSF barrier of the Choroid Plexus and reach the Arc from the 3rd ventricle by crossing the ependyma/tanycyte layer at the ventricle wall (1,2) ^6–11^. Hormones might enter the Arc through receptor-mediated transcytosis across BBB Arc vasculature (3)^6–11,37^. Fenestrated vessels of the ME can mediate entrance of hormones from the blood into the ME parenchyma, and then tanycytes lining the 3rd ventricle might facilitate the transport of hormones into the Arc (4,2)^6–11^. The pathways described above explain relatively slow or indirect mechanisms for hormonal exposure. Finally, ventro-medial (vm) Arc vessel bordering the ME might also be fenestrated^13^ (5).

**Extended Data Fig. 2.**
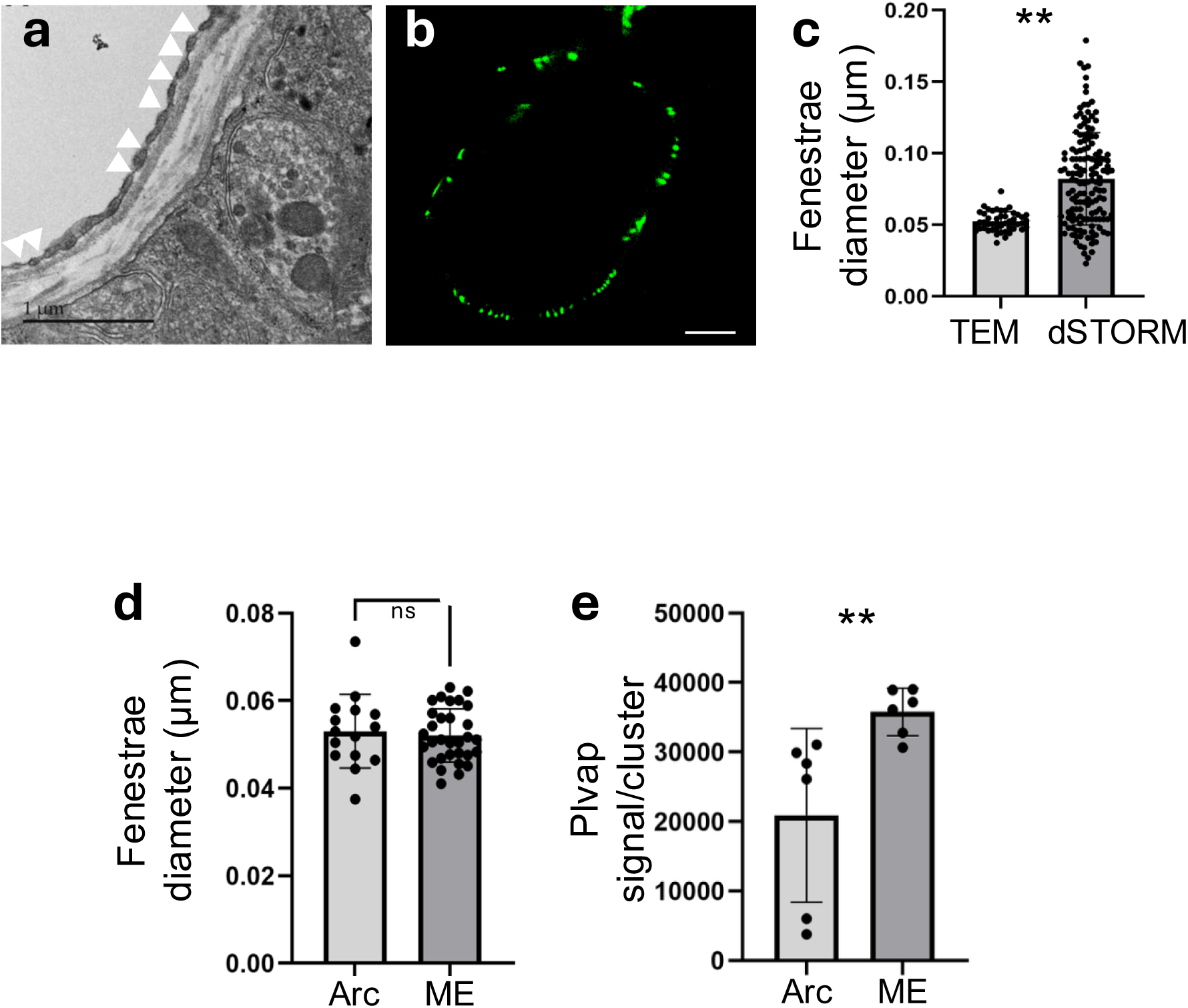
Ultrastructural dimensions of arcuate nucleus fenestrae correspond to nanoscale Plvap clusters detected by dSTORM. **a**, Representative TEM image of a fenestrated capillary showing endothelial fenestrae (arrowheads) as transcellular pores traversing attenuated regions of the endothelial cytoplasm. **b,** Representative dSTORM image of Plvap-labeled Arc capillaries revealing discrete nanoscale Plvap-positive clusters along the endothelial circumference. Each cluster serve as a proxy of a fenestrated domain. Scale bar, 1 μm. **c,** Comparison of Plvap cluster diameter measured by dSTORM with endothelial fenestra diameter measured by TEM, demonstrating close correspondence between the two measurements, in which each dSTORM cluster has the average size of 1.5 TEM fenestra (n ≈ 100 Plvap clusters/fenestrae per group; unpaired two-tailed Student’s t-test). ME and Arc capillaries have indistinguishable fenestrae diameter, but Plvap signal/cluster was higher at the ME. d, Quantification of endothelial fenestra diameter measured by TEM in capillaries from the median eminence (ME) and arcuate nucleus (Arc), demonstrating comparable fenestral dimensions between the two regions (n = 20 capillaries per group; unpaired two-tailed Student’s t-test). e, Quantification of Plvap signals per cluster measured by dSTORM in Arc and ME capillaries. Each data point represents one Plvap cluster average in mice capillaries (n=6). Data was collected from five mice. P < 0.01; two-tailed Mann–Whitney U-test.

**Extended Data Fig. 3.**
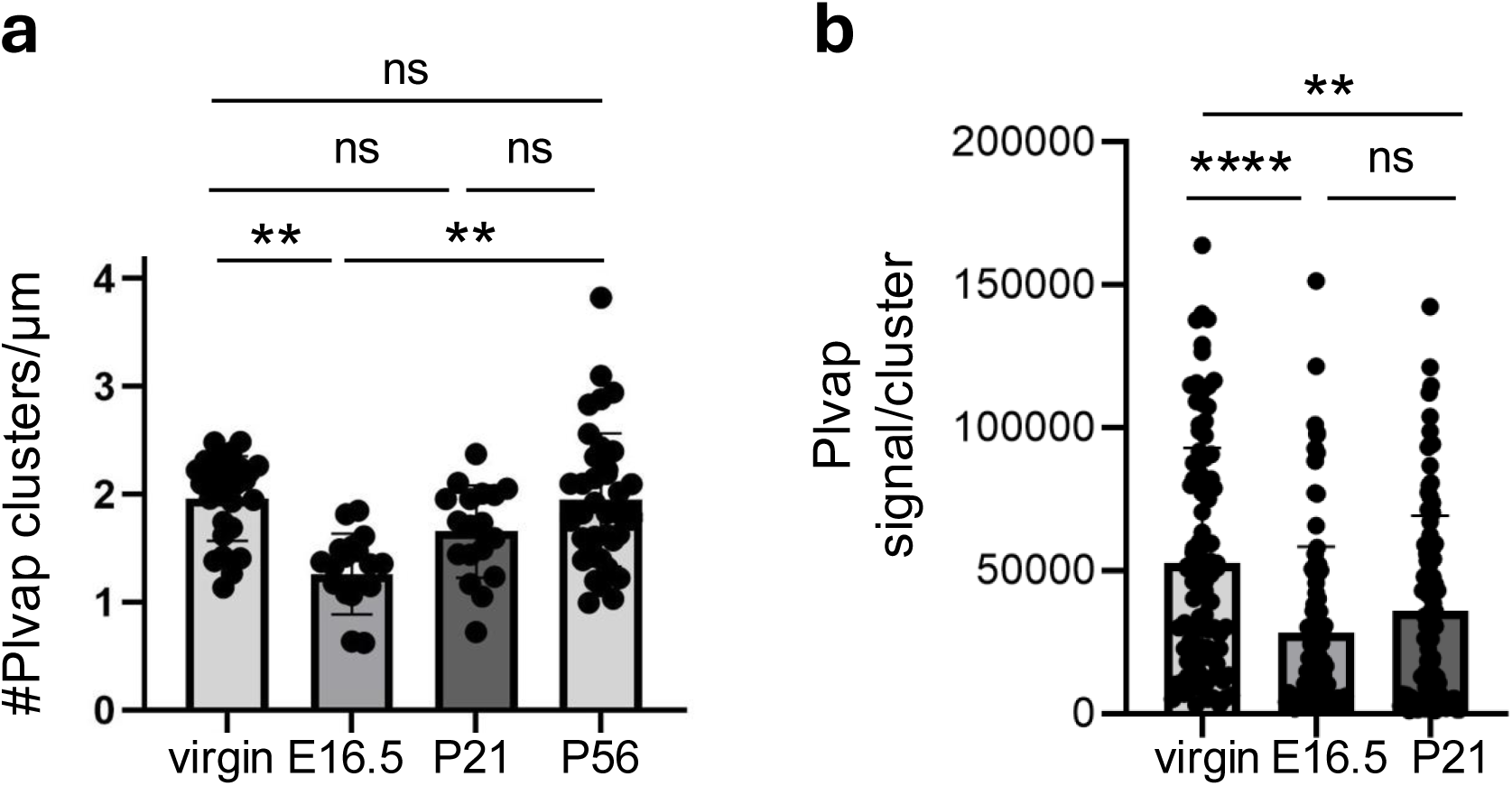
Remodeling of Arc fenestrations during pregnancy. **a**, Quantification of Plvap cluster density within Arc capillaries from dSTORM imaging data. Each data point represents one capillary (n ≈ 40 capillaries collected from 5 mice per group). **b,** Quantification of Plvap signal/cluster within Arc capillaries from dSTORM imaging data. Each data point represents one cluster (n ≈ 30 capillaries collected from 5 mice per group). ns, not significant; **P < 0.01, ****P < 0.0001; one-way ANOVA followed by Tukey’s multiple comparisons post-hoc test.

**Extended Data Fig. 4.**
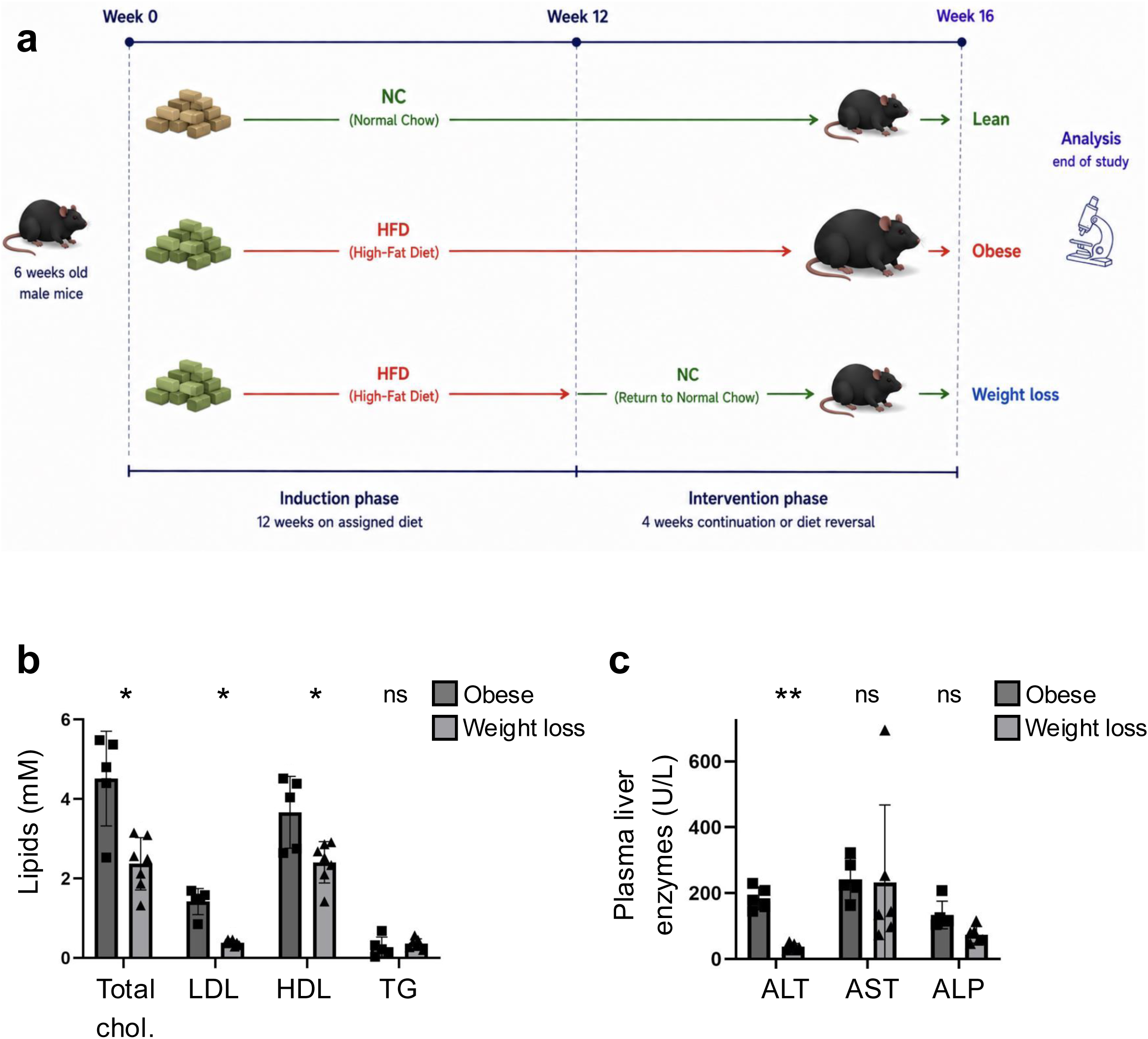
Remodeling of Arc fenestrations during diet-induced obesity. **a**, Experimental design of the diet-induced obesity and weight-loss model. Male mice (6 weeks old) were randomly assigned to one of three experimental groups: Lean, maintained on normal chow (NC) for 16 weeks; Obese, maintained on a high-fat diet (HFD) for 16 weeks; and Weight Loss, maintained on HFD for 12 weeks followed by 4 weeks of normal chow. At week 16, animals were euthanized for tissue collection and analysis, including TEM, dSTORM, serum biochemistry, and leptin transport assays. **b-c,** Levels of liver enzymes (ALT, AST, and ALP) and lipids (total cholesterol, LDL, HDL, and triglycerides) were mostly reduced in the weight-loss group compared to obese mice. Each data point represents one mouse (n = 5 mice per group) *P<0.05, **P<0.01; one-way ANOVA followed by Tukey’s multiple comparisons post-hoc test. Extended text related to main Fig. 2 A recent study investigated the receptor-mediated transcytosis pathway at the BBB by genetically eliminating Leptin Receptor in BBB endothelial cells37. This perturbation had no effect on mice until they were fed a high fat diet, causing them to gain more weight and faster. A possible explanation for this result would be that on regular chow, leptin signal is able to reach the Arc through fenestra, overriding receptor-mediated transcytosis. Once mice gain weight, the fenestra may close, making slower pathways such as Leptin-Receptor mediated transcytosis more central to leptin signaling. The dynamical aspects of hormonal resistance and the function of temporal scales for their signals to reach the Arc during pregnancy and obesity require further research.

**Extended Data Fig. 5.**
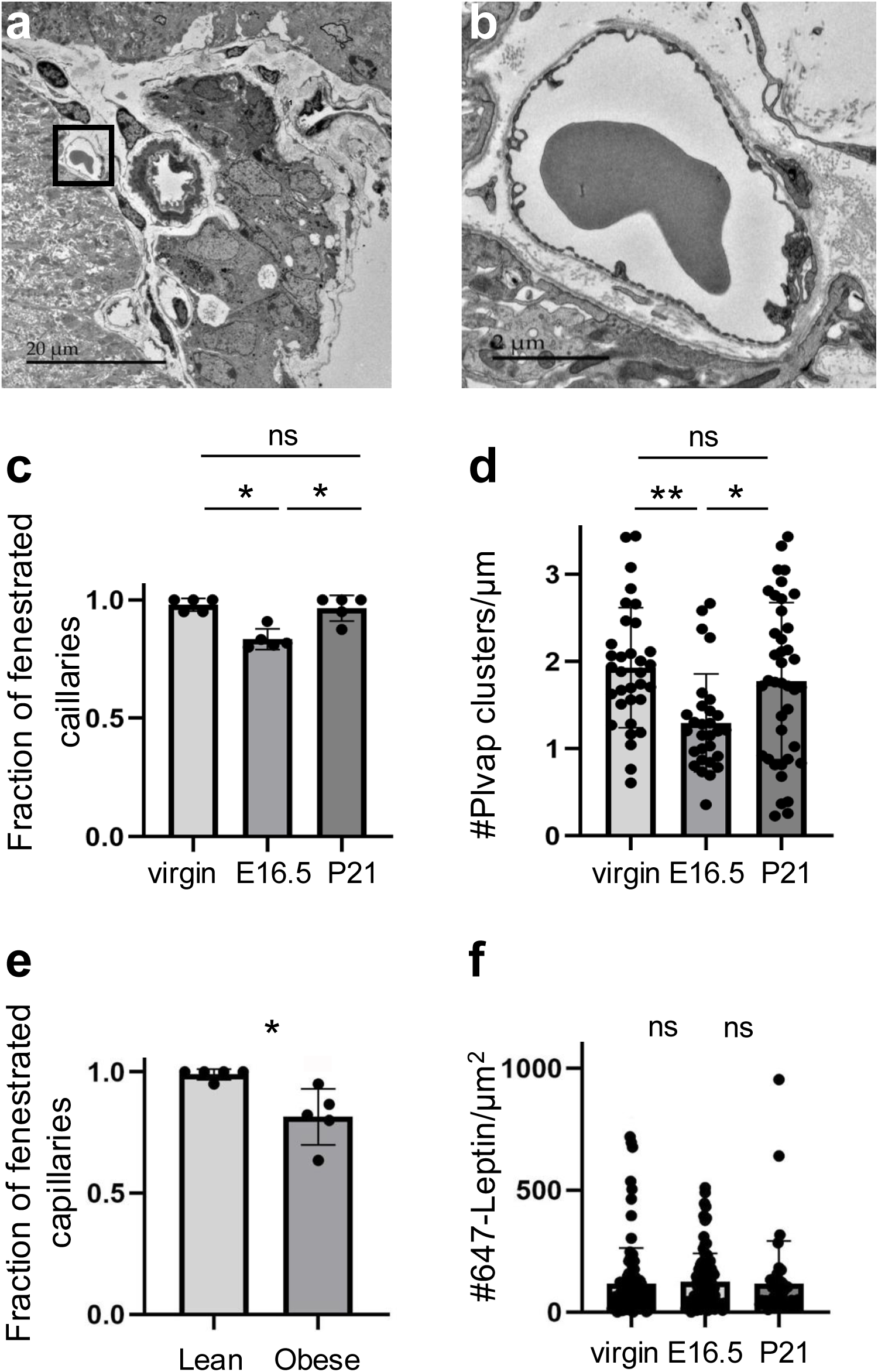
Changes in ME fenestrations during pregnancy and diet-induced obesity. **a-b**, Representative TEM images of median eminence (ME) capillaries from a virgin mouse and an E16.5 pregnant dam. Scale bar = 1 μm. **c,** Quantification of the proportion of fenestrated capillaries in the ME in virgin, E16.5 pregnant, and 21 days postpartum, based on TEM analysis. Each data point represents the average proportion of fenestrated capillary cross-sections quantified from a single mouse (n = 5 mice per group). **d,** Quantification of endothelial fenestra density within fenestrated ME capillaries measured by TEM. Each data point represents one fenestrated capillary (n = 30 capillaries collected from 5 mice). **e,** Quantification of the proportion of fenestrated capillaries in the ME by TEM in Lean and Obese mice. Each data point represents one mouse (n = 5 mice per group). **f,** Quantification of leptin fluorescence in the median eminence across pregnancy course. Each data point represents one fenestrated capillary (n ≈ 50 capillaries collected from 5 mice per group). *P < 0.05, \**P < 0.01; one-way ANOVA followed by Tukey’s multiple comparisons post hoc test*.

**Extended Data Fig. 6.**
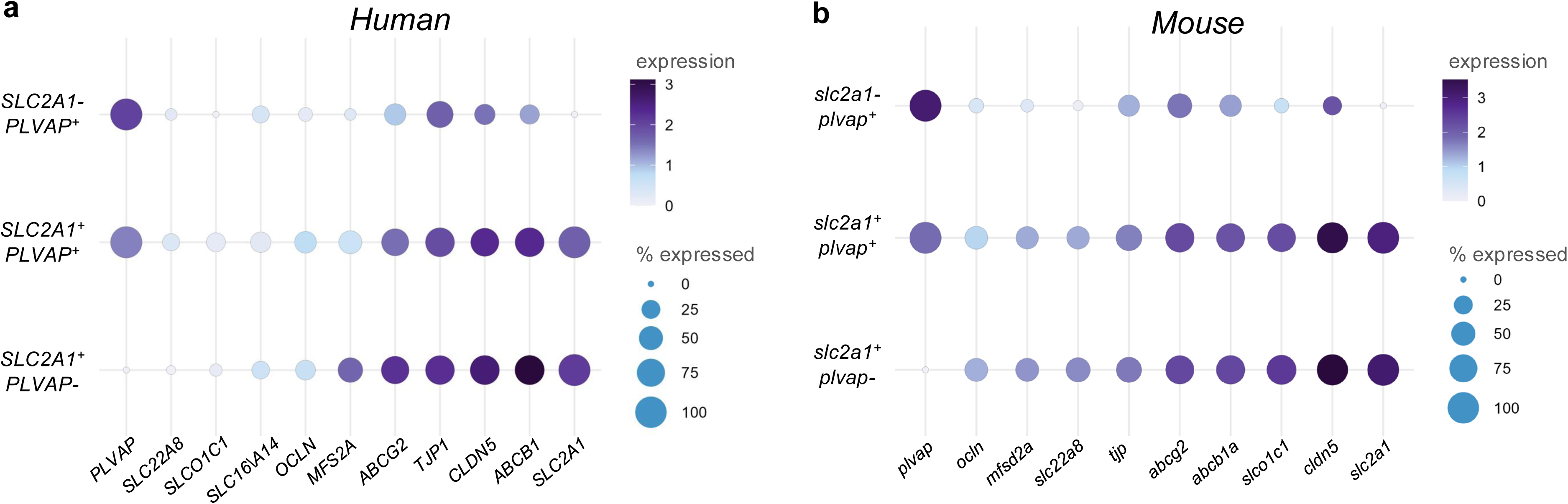
Unique BBB-permeable hybrid ECs co-express *PLVAP* with central BBB-associated genes. **a-b** Dot plot analyses of published human (a) and mouse (b) hypothalamic single-cell RNA sequencing datasets identifying endothelial populations expressing PLVAP, SLC2A1 (GLUT1), or both markers. Dot size represents the percentage of cells expressing each marker, and color intensity indicates normalized gene expression. Extended text related to Fig. 6 The hybrid ECs identified by these analysesd, are not likely to be mere technical doublets of scRNAseq. The data passed rigorous quality control, and these cells were identified in two independent datasets in mice and another dataset in humans. Second, Plavp+Glut1+ cells were directly observed using dSTORM in mice at the protein level (main Fig. 3a).

**Extended Data Fig. 7.**
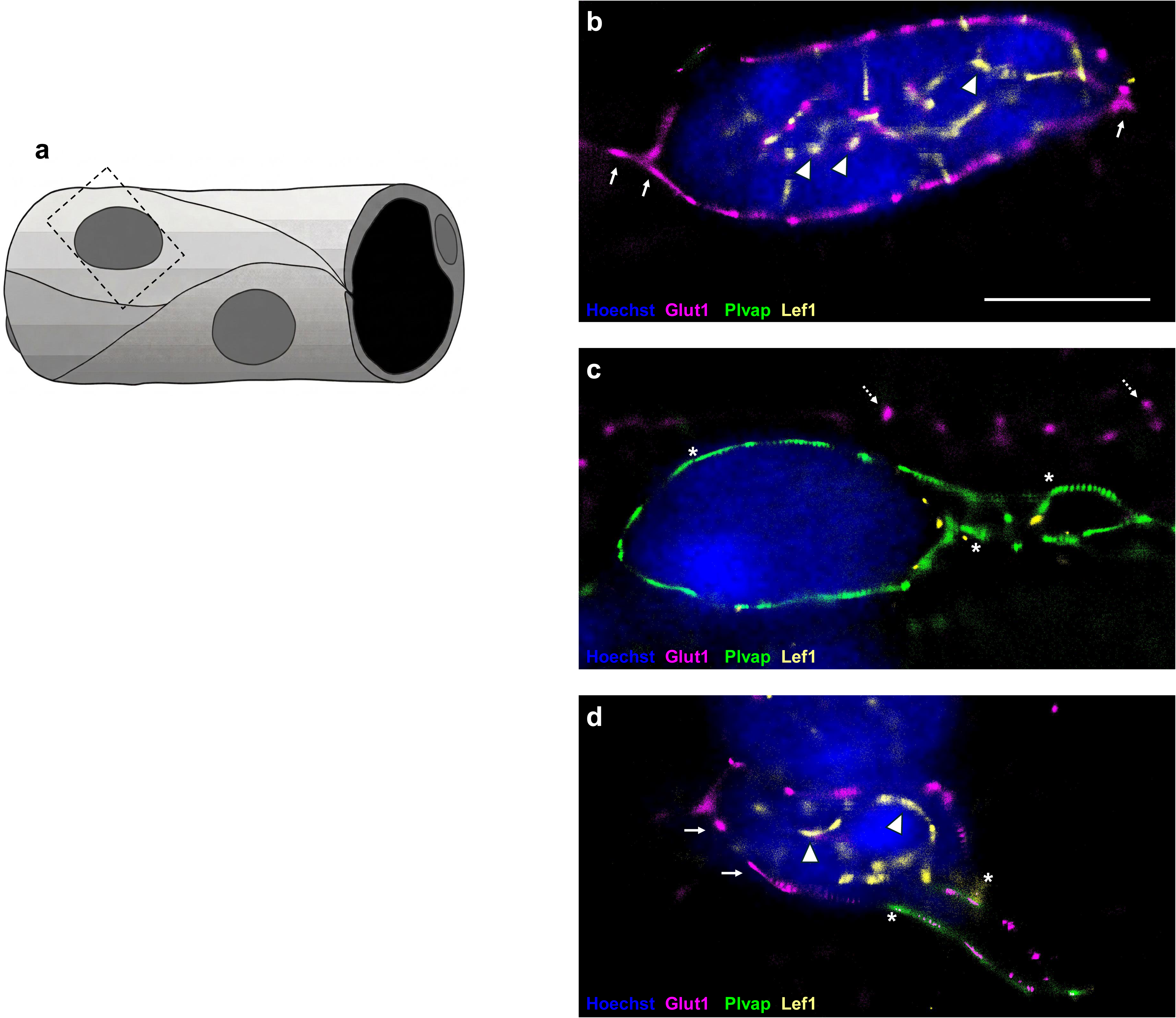
Wnt signaling is associated with endothelial populations. **a,** Illustration of dSTORM imaging approach - capillaries were identified by Plvap and Glut1 signals, and Nuclei by Hoechst signals in wide-field mode. Imaging angle was adjusted to include a wide nucleus section in wide-field mode. Hoechst shows at wide-field view, while Glut1, Plvap and Lef1 are imaged in 2D-dSTORM mode, resulting in cell membrane signals immediately around the nucleus with some signals that appear inside are probably below or above the nucleus plane. **b-d,** Representative dSTORM images of Cortex (b), ME (c), and Arc (d) EC nuclei. Endothelial Glut1 signals are clear at the cortex and Arc (arrows in b and d). Glut signal at the ME is probably astrocytic (dashed arrows in c). Plvap signals are clear at the ME and Arc (asterisk c and d). Nuclear Lef1 signals are clear at the cortex and Arc (arrow heads in b and d) but not at the ME. SMLM imaging allow accurate quantification of Lef1 signals (see main Fig 3h). Scale bar = 5 μm.

**Extended Data Fig. 8.**
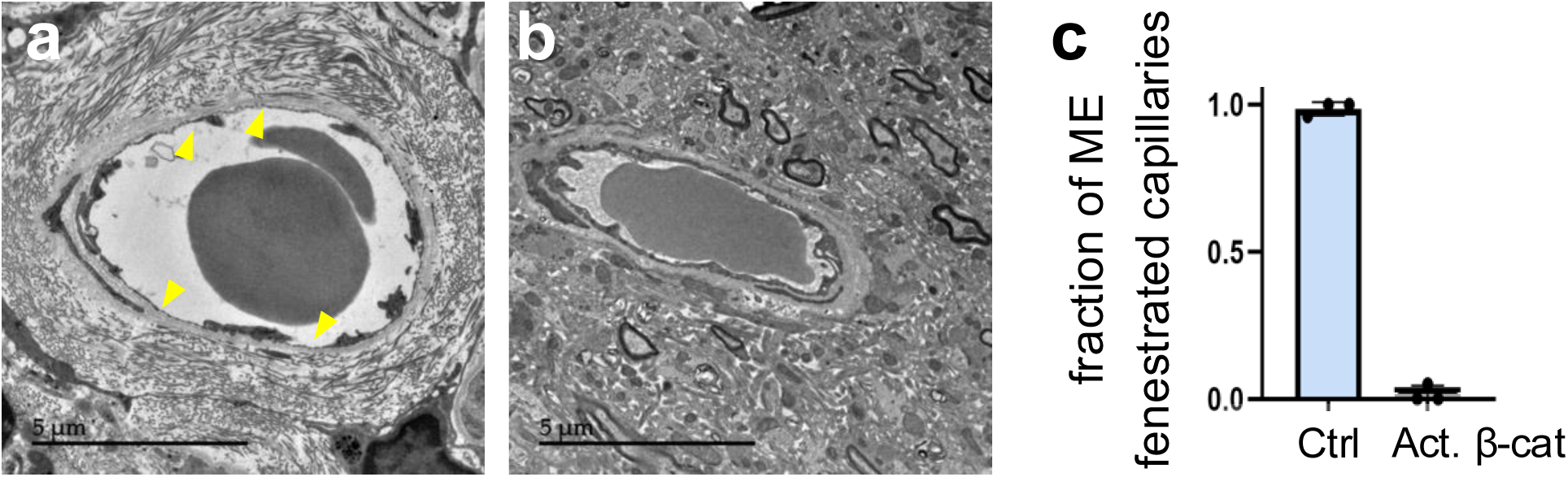
β-catenin activation suppresses fenestration in hypothalamic capillaries. **a-b**, Representative TEM images of ME capillaries from control (a) and β-catenin-activated mice (b). Yellow arrowheads indicate endothelial fenestrae. Scale bars = 5 μm. **c,** Quantification of the proportion of fenestrated capillaries in the ME of control and β-catenin-activated mice. Each data point represents one mouse (n = 3 mice per group).

**Extended Data Fig. 9.**
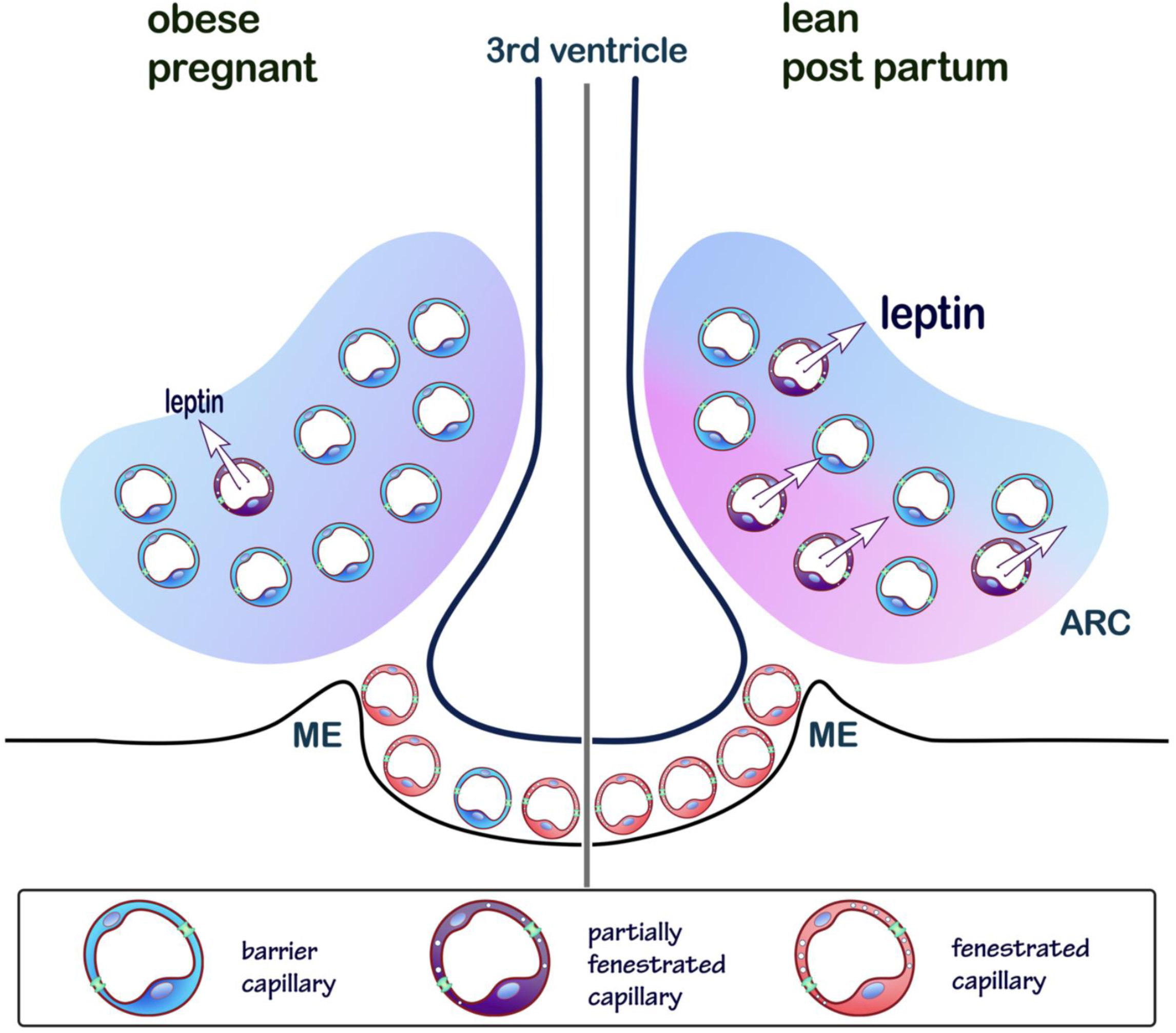

**Extended Data Fig. 10.**
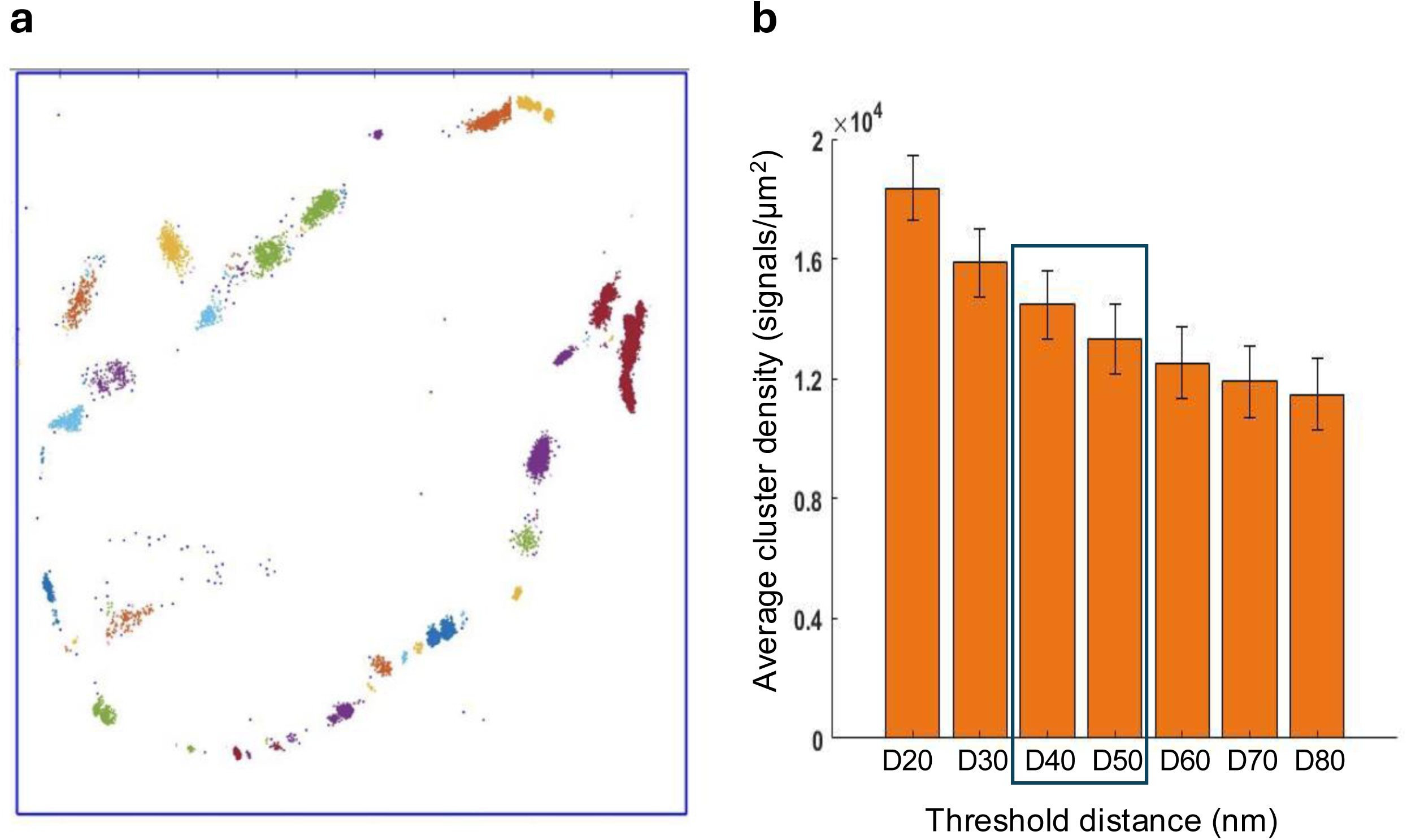
Super-resolution analysis methodology of PLVAP clusters. A simulation of Plvap clustering analysis in a representative capillary; each cluster is labeled in one color. **b** Plvap signal density was quantified across multiple segmentation thresholds (D20–D80), demonstrating no statistical differences between thresholds and supporting the selection of D50, corresponding to the average endothelial fenestra diameter determined by TEM.

